# Uncovering the cell type specificity of blood sample derived gene signatures using RNA expression data

**DOI:** 10.1101/684159

**Authors:** Mikhail Pomaznoy, Brendan Ha, Bjoern Peters

**Affiliations:** Division of Vaccine Discovery, La Jolla Institute for Immunology, La Jolla, CA; Department of Medicine, University of California San Diego, La Jolla, CA, United States

## Abstract

Analysis of transcriptomic data derived from blood samples is complicated by the complex mixture of cell types such samples contain. Transcriptomic signatures derived from such samples are often driven by a particular cell lineage within the mixture. Identifying this most contributing lineage can help to provide a biological interpretation of the signature. We created a web application CellTypeScore which quantifies and visually represents the expression level of signature genes in common blood cell types. This is done by constructing an interactive stacked bar plot with the bars representing expression of genes across blood cell types. Summed scores serve as a measure of how highly the combined signature is expressed in different cell types. An online version of the application can be found at https://tools.dice-database.org/celltypescore/.

## Background

Immunology research has widely adopted transcriptomic profiling which results in gene signatures characterizing a particular immune perturbation. For example, a signature can be comprised of genes differentially expressed between infected and uninfected individuals, or it can be a list of genes co-expressed longitudinally after vaccination event, etc. Starting material for such studies is often whole blood or PBMCs which is a complex mixture of various cell populations. For biological interpretation of a signature it is crucial to determine if the signature can be attributed to a particular cell type or a lineage of immune cells. Sometimes a signature may be explained by a frequency change of one of the cell types or a transcriptional program shift within only one lineage.

To facilitate the process of establishing relation of signatures to cell types we used an advantage of recently developed DICE database^1^ resource. This database contains uniformly processed high quality expression data of common cell types found in peripheral blood. We used this expression data to create a web application CellTypeScore which creates an interactive stacked bar plot. The plot represents the sum of expression of signature genes in different cell types available in DICE-DB. It allows to estimate which cell type is the most likely to contribute to the signature and explore genes and corresponding cell types through the DICE database resource.

## Results

### User workflow

User input consists of a list of human genes uploaded as a plain text file or pasted into a textbox. After processing the input application, the tool outputs a page as shown in in Fig. 1. The output consists of an SVG image with a stacked bar plot and control switches affecting the plot. X-axis corresponds to cell types and Y-axis represents expression score as a sum of TPM values of every gene in the signature in a particular cell type.

**Figure 1.**
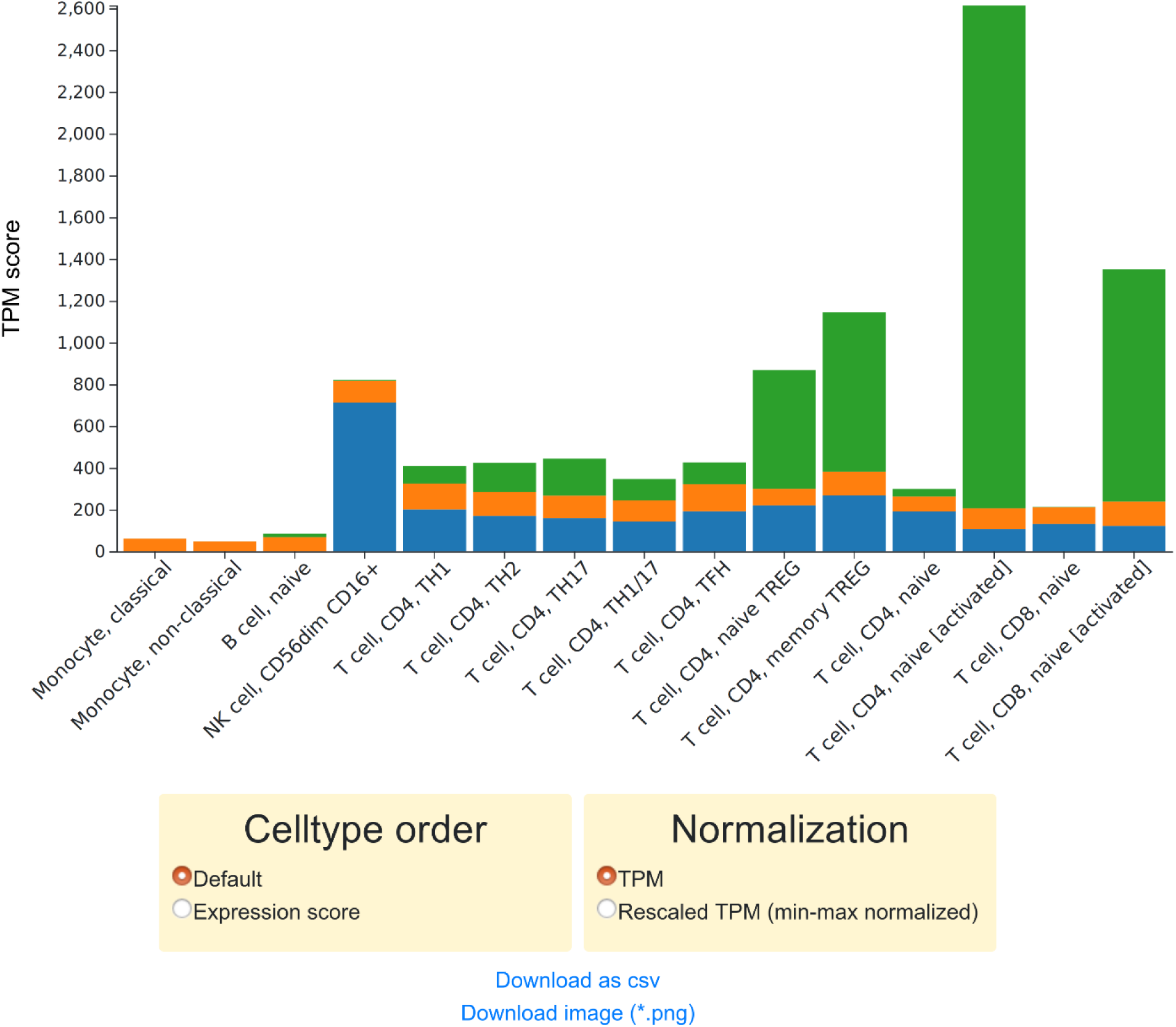
Stacked bar plot constructed by CellTypeScore application. A sample signature of three genes CD3Z (blue), NFKB3 (orange), IL2RA (green) was used as input.

The first control switch allows the user to change the order of cell types in the X-axis. Available options are:

- **Default order.** This is a predefined order that displays related cell types next to each other.
- **Expression score.** This is a data driven order, ordering cell types from highest to lowest expression score.

The second control switch allows the user to choose normalization of expression values:

- **TPM** simply displays the sum of raw TPM values of the input genes.
- **Rescaled TPM** utilizes a min-max normalization that sets expression of every gene normalized to a range of [0, 1] across all the cell types. This makes contributions of genes to the score more equal. It is especially useful when one gene is very highly expressed compared to the others, because in this situation overall score can be driven by a single outlier gene.

Additionally interfaces allows user to download the result in a .CSV format containing information about how the input gene names were converted to machine readable IDs and expression of each input gene in across the cell types available in DICE-DB. Additionally .PNG image of the plot can be downloaded for illustrative purposes.

Importantly the gene boxes are interactive and display information about the underlying gene when hovered above. It also allows to easily access DICE database page for a particular gene entry.

### Application to known signatures

For validation purposes we wanted to test our application on lists of known cell type-specific genes derived on different platforms than the one used for generation of DICE database. We extracted genes distinguishing hematopoietic cells from the study by Newman et al^2^, where the authors combined several previous studies in which blood populations were profiled using microarrays to derive specific markers. These cell type-specific markers were then used for estimating cell type frequencies from expression data of a mixed sample. We picked those genes which are unique for a single cell type only and performed CellTypeScore analysis on those genes. Results of min-max normalized expression scores are shown in Figure 2.

**Figure 2.**
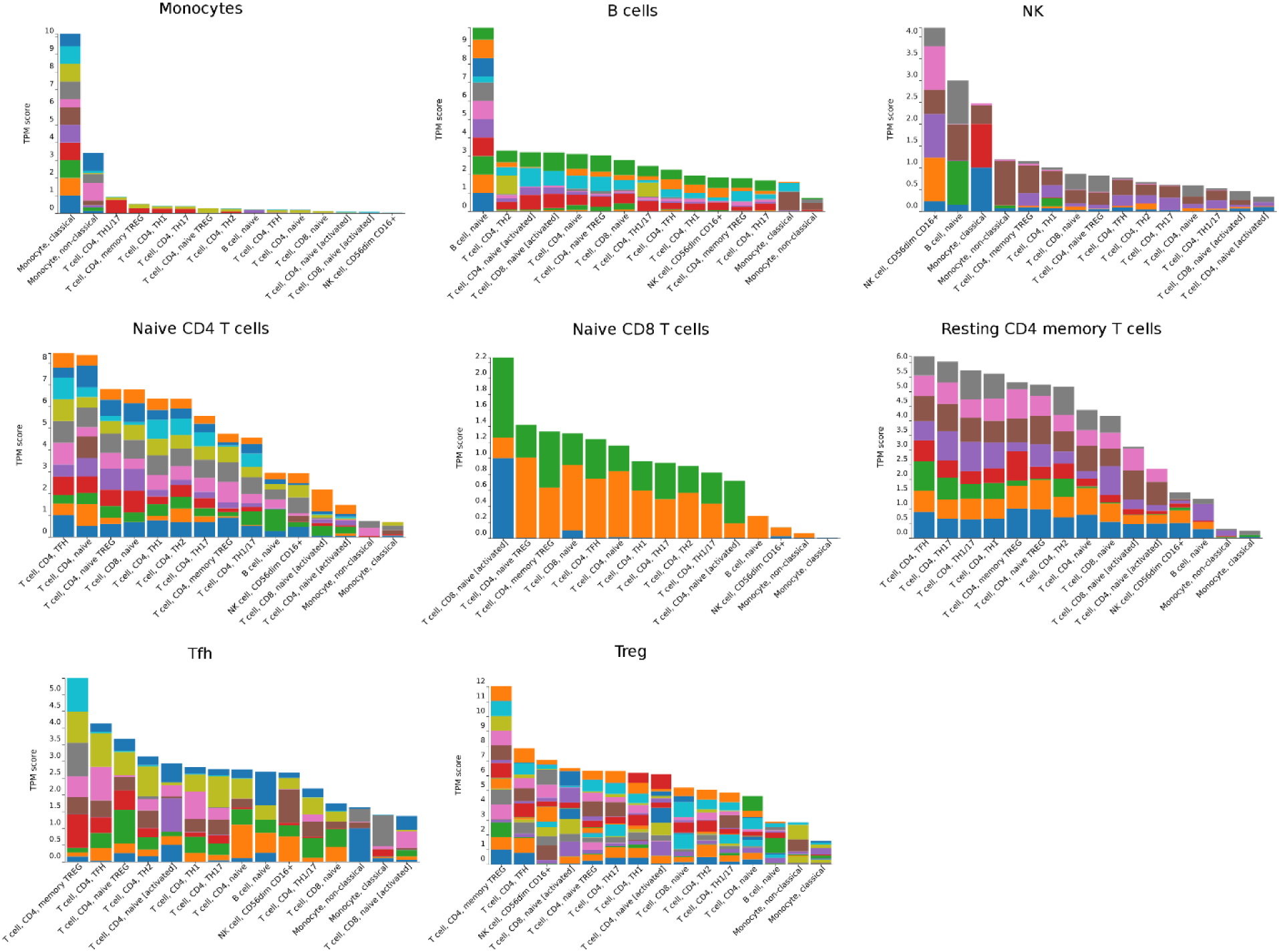
CellTypeScore results for cell-type specific genes obtained in CIBERSORT. Titles of subplots indicate genes specific for which cell type were captured.

We observed consistent and expected scores for monocytes, B cells and NK cells (Fig. 1, top row), i.e. marker genes from this cell types have the highest score against the DICE-DB counterpart. Treg-specific genes were also identified as expected. Resting CD4 memory concordantly with the definition had highest scores against different CD4 subsets (T_FH_, T_H_17, T_H_1 etc.). Naive CD8 T cell-specific genes did not match with corresponding DICE-DB CD8 T cells. But there were only three of these genes extracted from Newman et al, making overall signature assignment ambiguous. Naive CD4 T cells had actual DICE-DB counterpart as the second highest score while demonstrating the highest score against T_FH_. Interestingly, T_FH_-specific genes had the highest score against Treg memory cells while having the second highest score against actual T_FH_ from DICE-DB.

We further wanted to validate our approach on transcriptomic signatures of diseases. We specifically investigated signatures differentiating active tuberculosis (TB) patients from healthy or latently infected individuals. There are several studies on this subject because differentiating active tuberculosis from latent infection is an important diagnostic problem (e.g. see^3,4^). We extracted signature distinguishing patients with active TB from latently infected or healthy individuals from study by Kaforou et al. performed on cohorts from South Africa and Malawi^5^ and a signature from a study by Walter et al performed on US patients cohort^6^. We analyzed separately genes up- or down-regulated in active TB patients from both studies. Resulting output of CellTypeScore is shown in Figure 3. Despite significant differences in gene content of the signatures, the results of the CellTypeScore look rather similar. Genes up-regulated in active TB patients possess high score for monocytes (classical in particular). Genes down-regulated in active TB patients on the opposite have very low score for monocytes and are mainly expressed in B cells for the signature from Kaforou et al and in T cells for the signature from Walter et al.

**Figure 3.**
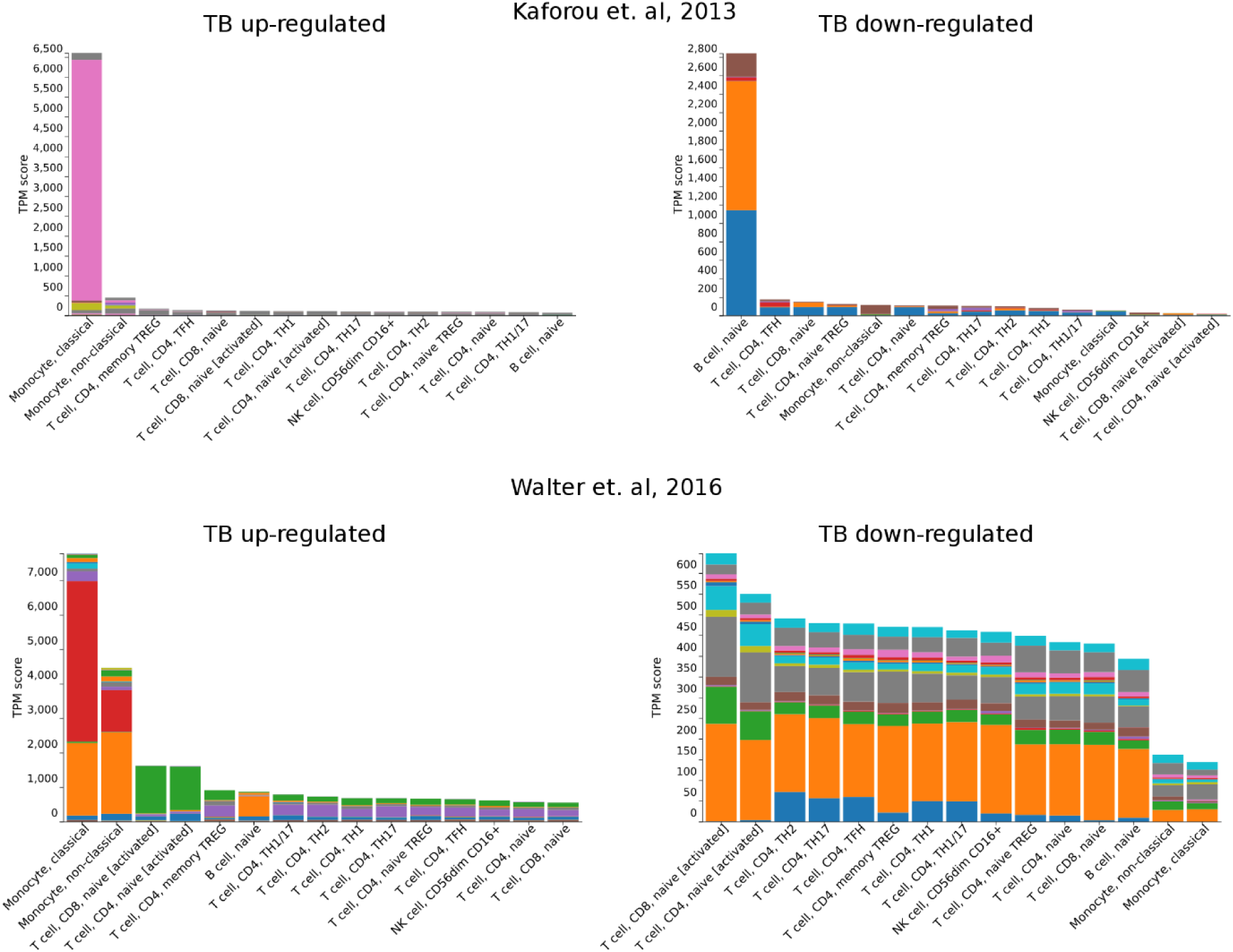
Application of CellTypeScore to signatures of active TB.

## Implementation

### Back-end

Python Django is used as a framework for back-end implementation.

Firstly, user input is converted to standardized gene ID. For this purpose we internally use a gene name resolver which compiles canonic gene symbols, unique gene IDs from widely used resources, and manually curated immunologist-preferred labels into a single source to resolve gene queries. The gene name resolver takes a set of gene names as input and attempts to resolve them to canonical gene symbols which are also used in the DICE dataset. Potential gene names are categorized into the following in order of decreasing precedence: canonical gene names, synonyms, or immunologist preferred labels. Potential gene names are then scored based on string distance and assigned the best scoring gene as the resolved gene.

The resolved genes are then used to query expression data from the DICE dataset which will be sent to the client for visualization.

### Front-end

The bar plot is generated using D3 JavaScript library^7^.

## Discussion

Transcriptomic signatures derived from immunology studies are often hard to interpret if the starting biological material are whole blood or PBMC samples. In some cases only a certain cell type lineage contributes to the signature. In other cases signature can be driven by major changes in cell type composition. This uncertainty complicates biological interpretation of the data. Here we present an application which scores a given signature against blood cell types helping to identify cell type(s) contributing the most.

We applied our approach to gene distinguishing hematopoietic cell phenotypes derived from the study Newman et al^2^. We found that genes specific for broad and relatively homogeneous cell types like monocytes or B cells are clearly identified using CellTypeScore. On the other hand distinguishing separate T cell populations is a harder task. This is expected due to diverse nature of T cells and relative proximity of T cell subsets to each other. This indicates limitations on resolution of certain signatures up to T cells subsets. Nevertheless, assigning a signature to a broad classes of cell populations is more robust.

We further applied CellTypeScore to signatures differentiating whole blood of active TB patients from healthy people. We found that up-regulated genes had high score against monocytes. On the contrary genes down-regulated in active TB had very low score against monocytes and were expressed highest in B or T cells. It highlights the possibility that these signatures can be explained by a change in frequency of monocytes. However DICE-DB is currently lacking data for such myeloid cell types as neutrophils, DCs etc. These cells are more similar to monocytes than to lymphoid cells and could be another likely source of these TB signatures. For example, in a study by Berry et. al^8^ authors conclude that active TB signature is neutrophil-driven. However, increase in monocyte/lymphocyte count ratio is a known marker of active tuberculosis infection^9^ and monocyte frequency could very likely drive signatures of active TB.

In conclusion we believe that our approach is a useful tool for blood transcriptomic studies and can be further improved as transcriptional data for additional cell types is getting available.

